# Characterization of Thylakoid Division Using Chloroplast Dividing Mutants in *Arabidopsis*

**DOI:** 10.1101/200667

**Authors:** Jonathan Ho, Warren Kwan, Vivian Li, Steven M. Theg

**Author notes:** **Summary:** Chloroplasts unable to divide possess giant thylakoids suggesting that thylakoids do not possess independent division machinery. **Author Contributions:** JH and SMT designed experiments. JH, WK and VL performed the experiments and all authors analyzed the data. JH and SMT wrote the article and all authors approved it. **Funding information:** This work was funded by the Division of Chemical Sciences, Geosciences, and Biosciences, Office of Basic Energy Sciences of the US Department of Energy through Grant DE-FG02-03ER15405.

## Abstract

Chloroplasts are double membrane bound organelles that are found in plants and algae. Their division requires a number of proteins to assemble into rings along the center of the organelle and to constrict in synchrony. Chloroplasts possess a third membrane system, the thylakoids, which house the majority of proteins responsible for the light-dependent reactions. The mechanism that allows chloroplasts to sort out and separate the intricate thylakoid membrane structures during organelle division remain unknown. By characterizing the sizes of thylakoids found in a number of different chloroplast division mutants in *Arabidopsis*, we show that thylakoids do not divide independently of the chloroplast division cycle. More specifically, we show that thylakoid division requires the formation of both the inner and the outer contractile rings of the chloroplast.

## Introduction

Chloroplasts are bound by outer and inner envelope membranes which enclose an additional compartment called the thylakoid. Specialized to capture the light energy available in sunlight and convert it into ATP and NADPH, thylakoids are responsible for the production of most of the oxygen in the Earth’s atmosphere.

Structurally, thylakoids appear as an intricate network of columns of stacked discs, termed grana, interconnected by unstacked regions called stromal lamellae. This architectural complexity is further highlighted by the location of different proteins complexes within the different thylakoid regions. Photosystem II and the light-harvesting complex II trimer are found in the granal regions of the thylakoid, whereas the photosystem I - light harvesting complex I supercomplex and the ATP synthetase are found in the stromal lamellae; the cytochrome b6/f complex is located in both (Boardman et al., 1966; Boekema et al., 2000; van Roon et al., 2000; Daum et al., 2010). Spectroscopic studies, electron microscopy (Heslopharrison, 1963; Paolillo and Falk, 1966; Schoenknecht et al., 1990) and more recently, electron tomography (Shimoni et al., 2005; Daum et al., 2010; Austin and Staehelin, 2011) have shown that the thylakoid system within a single chloroplast consists of a single vesicle with its membrane intricately folded upon itself. A number of proteins, such as Thf1, VIPP1, FtsZ and FZL were shown to be essential for the development and maintenance of thylakoid structure (Kroll et al., 2001; Wang et al., 2004; Gao et al., 2006; El-Kafafi et al., 2008; Karamoko et al., 2011; Lo and Theg, 2012). The mechanism that allows chloroplasts to sort and separate the intricate structures of the thylakoid membrane during organelle division remain unknown.

Chloroplasts division requires a multitude of proteins to assemble into rings along the center of the organelle and to constrict in synchrony (Maple-Grodem and Raynaud, 2014; Osteryoung and Pyke, 2014). The first ring that forms involves the stromal protein FtsZ interacting with Arc6, a protein found in the inner envelope membrane (McAndrew et al., 2001; Vitha et al., 2003). Arc6 has also been shown to interact with PDV2 in the outer envelope, which along with PDV1, recruits DRP5 or Arc 5, a dynamin-related protein that forms a contractile ring along the cytoplasmic side of the chloroplast (Yoshida et al., 2006; Glynn et al., 2008). FtsZ1 and isoforms of FtsZ2 in *Arabidopsis* have been found to associate with the thylakoid membrane in a developmentally dependent manner (El-Kafafi et al., 2008; Karamoko et al., 2011). While this is consistent with the involvement of these proteins with thylakoid division, this remains a matter of speculation.

The separation of thylakoids during chloroplast division has been captured in electron micrographs by a number of different groups (Leech et al., 1981; Oross and Possingham, 1989; Robertson et al., 1996). Images show thylakoids dispersed throughout the chloroplast in the early stages of plastid division. As division proceeds, the number of thylakoid grana and stromal lamellae that extend across the length of the original plastid dwindles until a single segment spans the isthmus of the constricted plastid. At the final stages of plastid division, the thylakoid membrane along with one of the daughter organelles appear to be twisted so that the thylakoid membrane loses its longitudinal orientation (Robertson et al., 1996). Thylakoid division appears to precede the separation of the daughter organelles (Whatley, 1980). The mechanism of thylakoid division, and its reliance on the chloroplast division cycle remain unknown. In this study we dissect the dependence of thylakoid division on the chloroplast division machinery by comparing the sizes of thylakoids found in chloroplast division mutants that are arrested at different states of plastid division.

## Results

### Experimental theory and design

Wild type *Arabidopsis* plants possess ~ 120 chloroplasts per mesophyll cell, each with a single thylakoid membrane (Pyke and Leech, 1994). The composition of thylakoids from chloroplast mutants that are unable to divide is unknown. We first examined the effect of the lack of an inner contractile ring on the division of thylakoids. The chloroplast division mutant, *arc6*, has a premature stop codon near the amino-terminal region of the protein, thereby preventing the formation of an inner contractile ring and rendering the plastid incapable of division (Vitha et al., 2003). As a result, the mutant plant possess giant-sized chloroplasts (Pyke et al., 1994). We reasoned that there are two possible scenarios that can describe the fate of the thylakoids in *arc6* mutants. In the first, we assume that the thylakoids in the *arc6* chloroplasts are still capable of undergoing division independent of the chloroplast division cycle. As a result, the number and size of the thylakoids within the giant *arc6* chloroplasts will likely be similar to those found in wild-type cells. In the second scenario, we assume that the *arc6* thylakoids cannot divide independently of the chloroplast division cycle. Accordingly, the *arc6* chloroplasts would possess a single giant-sized thylakoid.

We sought to determine the relative sizes of the thylakoids in *arc6* and wild type chloroplasts by measuring the sensitivity of the ionic conductivity of the thylakoid membrane to the pore-forming ionophore gramicidin. This experiment, originally performed by (Schoenknecht et al., 1990), is based on the ability of a small number of gramicidin pores to short circuit the capacitance of a thylakoid membrane vesicle. Titration of the membrane capacitance with gramicidin will report on the size of the thylakoid electrical unit, with larger thylakoids being more readily short-circuited by a given concentration of ionophore than smaller thylakoids. The conductivity of the thylakoid membrane can be conveniently and non-invasively monitored via the well-characterized decay of the carotenoid electrochromic shift at 520 nm induced by a short pulse of light (Junge and Witt, 1968; Witt, 1979; Bailleul et al., 2011). By this technique it was determined that the size of the thylakoid electrical unit within chloroplasts corresponds to all the photosynthetically active membranes within each plastid; that is, there is essentially one thylakoid per chloroplast (Schoenknecht et al. 1990). This conclusion has been confirmed by TEM micrographs of serially sectioned plastids (Heslopharrison, 1963; Paolillo and Falk, 1966; Mustardy and Janossy, 1979), and more recently, by electron tomography (Nierzwicki-Bauer et al., 1983; Shimoni et al., 2005; Mustardy et al., 2008; Austin and Staehelin, 2011; Daum and Kuhlbrandt, 2011).

### The electrical unit in *arc6* chloroplasts is larger than that in wild-type chloroplasts

It would be expected that large thylakoid vesicles potentially formed in giant chloroplasts might be susceptible to disruption by shear forces applied during isolation. In order to minimize this possibility, protoplasts were made from both wild type and *arc6* plants and chloroplasts were subsequently isolated therefrom. Figure 1 shows that despite the expectation that the giant single chloroplasts present in the *arc6* mutants would be even more fragile than those in the wild type, they could be isolated intact from protoplasts by incubation in a carbonate-containing buffer. This is, to our knowledge, the first successful isolation of these giant chloroplasts with their envelopes intact.

**Figure 1.**
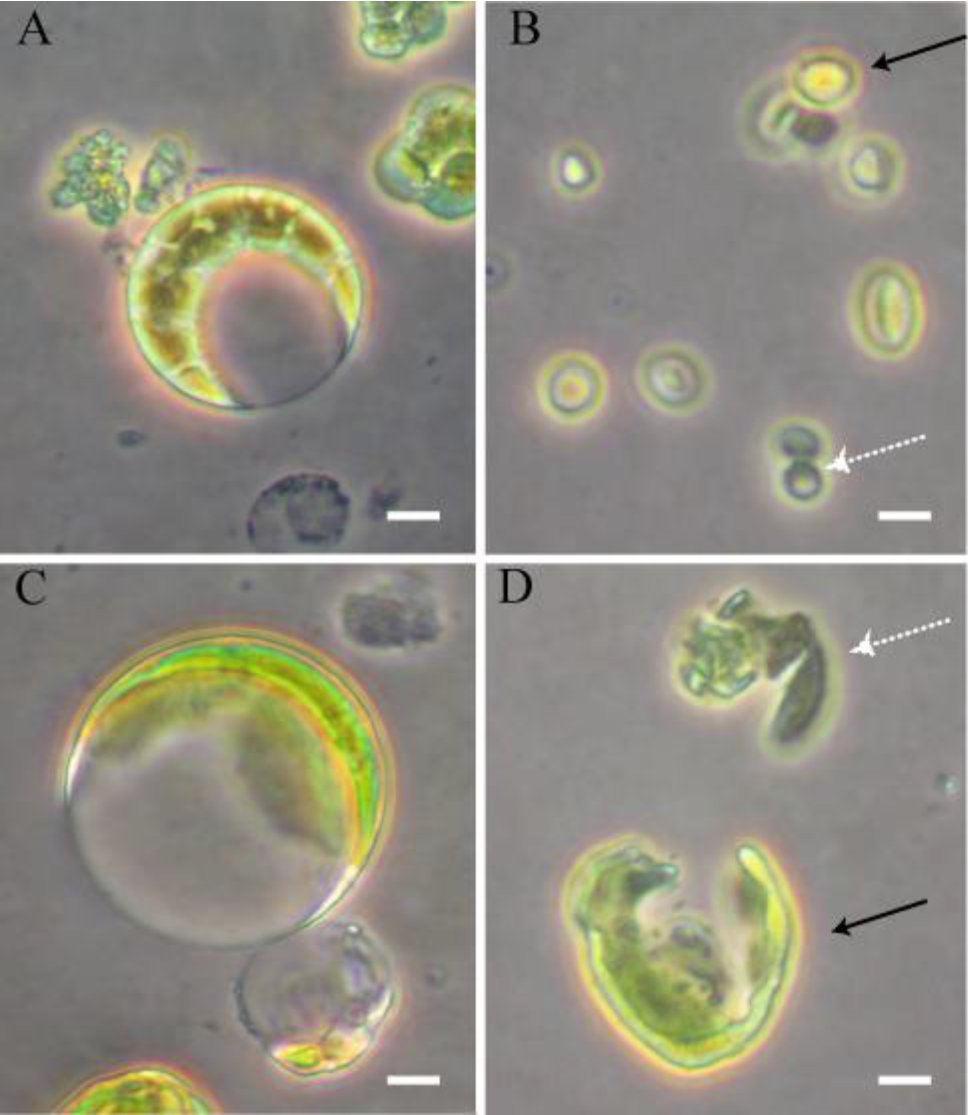
Isolation of intact chloroplasts. A) and C), Isolated protoplasts from wild type and *arc6 Arabidopsis* under a phase contrast microscope. B) and D), Wild type and *arc6* isolated chloroplasts. Intact chloroplasts possess a halo and are highlighted by the black arrow; broken chloroplasts are indicated by the white arrow. Scale bar corresponds to 10 μm.

We then probed the size of the thylakoid electrical units via the light-induced carotenoid electrochromic signal at 520 nm. After a 9 ms light pulse, the Δψ-indicating ΔA_520_ nm signal exhibited biphasic relaxation kinetics in both wild type and *arc6* plastids. *Arc6* chloroplasts displayed a faster Δψ relaxation rate than the wild type even in the absence of ionophore (Figure 2A-B). The addition of gramicidin to the chloroplasts resulted in an accelerated relaxation rate of the ΔA_520_ nm signal in both wild type and *arc6* samples (Figure 2A-B), with increasing concentrations resulting in faster decays. This was also conveniently manifested in the initial point recorded as the 9 ms actinic pulse was turned off. Since a 9 ms pulse is not a single turnover flash, the magnitude of the electric field measured at 9 ms results from competition between field generation by multiple reaction center excitations and decay by ion counter movement, the latter of which is accelerated by gramicidin. Accordingly, thylakoid membranes with increased ion permeability display a lower magnitude of the ΔA_520_ nm signal at 9 ms. Acceleration of the electrochromic shift decay can be clearly seen by plotting the magnitude of the 9 ms ΔA_520_ nm absorbance as a function of the logarithm of the gramicidin concentration (Figure 2C). This plot reveals the increased sensitivity of the thylakoid conductance to gramicidin exhibited by *arc6* chloroplasts over those from wild-type plants, and indicates that the thylakoids are larger in the *arc6* mutant.

**Figure 2.**
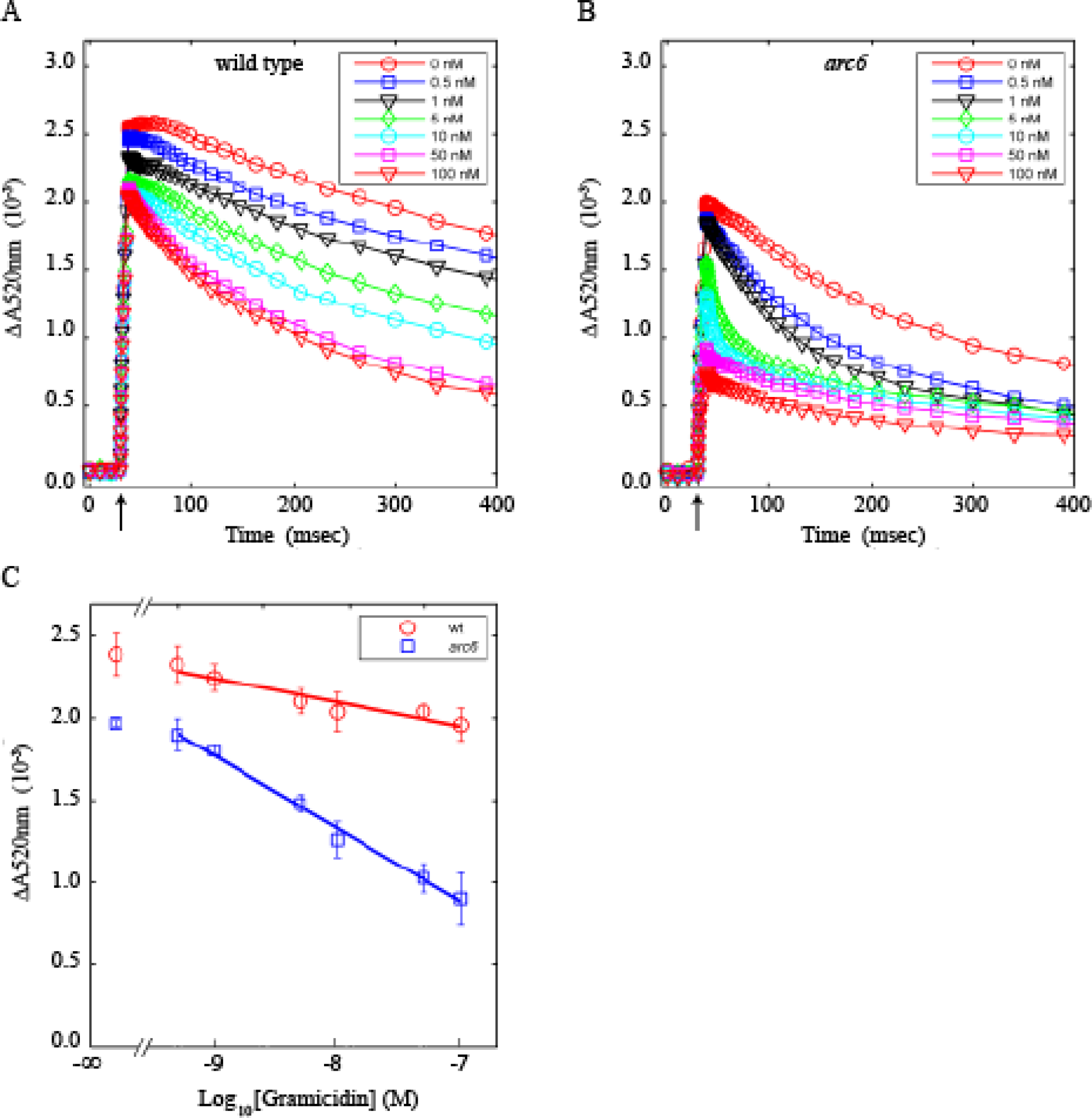
Thylakoid membrane conductivity measured by the electrochromic shift of carotenoids. Representative ΔA_520_ nm measurements of A) wild-type and B) *arc6* chloroplasts with increasing amounts of gramicidin; the arrow indicates the initiation of the 9 ms actinic flash. C) Plot of the first point of the ΔA_520_ nm signal after the 9 ms actinic flash vs. the log of gramicidin concentration. Bars indicate standard deviation of n=9 for each data point. The wild-type (red circles) and *arc6* (blue squares) data points fit by linear regression; the ΔA_520_ nm values for 0 nM gramicidin values are plotted to the left of the axis break and not utilized in the fit.

### *Arc6* chloroplasts possess giant thylakoids

When isolated protoplasts are placed in very low osmotic conditions, the plasma membrane and the chloroplast outer and inner envelope membranes break, resulting in the release of the thylakoids (Mercer, 1954; Weier et al., 1965; Hinnah and Wagner, 1998). However, thylakoid membranes are more resilient to osmotic pressure than the rest of the cell's membranes, and thylakoids swell to form blebs. We used this property of thylakoids to provide an independent measurement of the size of wild type and *arc6* thylakoids. In the confocal microscope, chlorophyll autofluorescence serves as a marker for thylakoid membranes (Figure 3). Figure 3B shows a representative cross section image of a bleb formed from *arc6* thylakoids, and it is apparent that it is significantly larger than those formed from wild-type thylakoids (Figure 3A). The diameter of the wild type bleb (Figure 3A) is ~ 12.8 ± 3.9 μm, whereas the *arc6* bleb (Figure 3B) diameter averages 21.7 ± 11.2 μm. While the diameter of a spherical object such as a bleb observed in a single cross section could be mistakenly underestimated by examining an image off of the equatorial plane, those images in Fig. 3 were produced after scanning back and forth through the z-plane to find the maximum observed diameters. Representative z-projections, in which all z-plane images are stacked one upon each other, confirm the dramatic difference in sizes between *arc6* and wild type blebs (Figure 4A-B).

**Figure 3.**
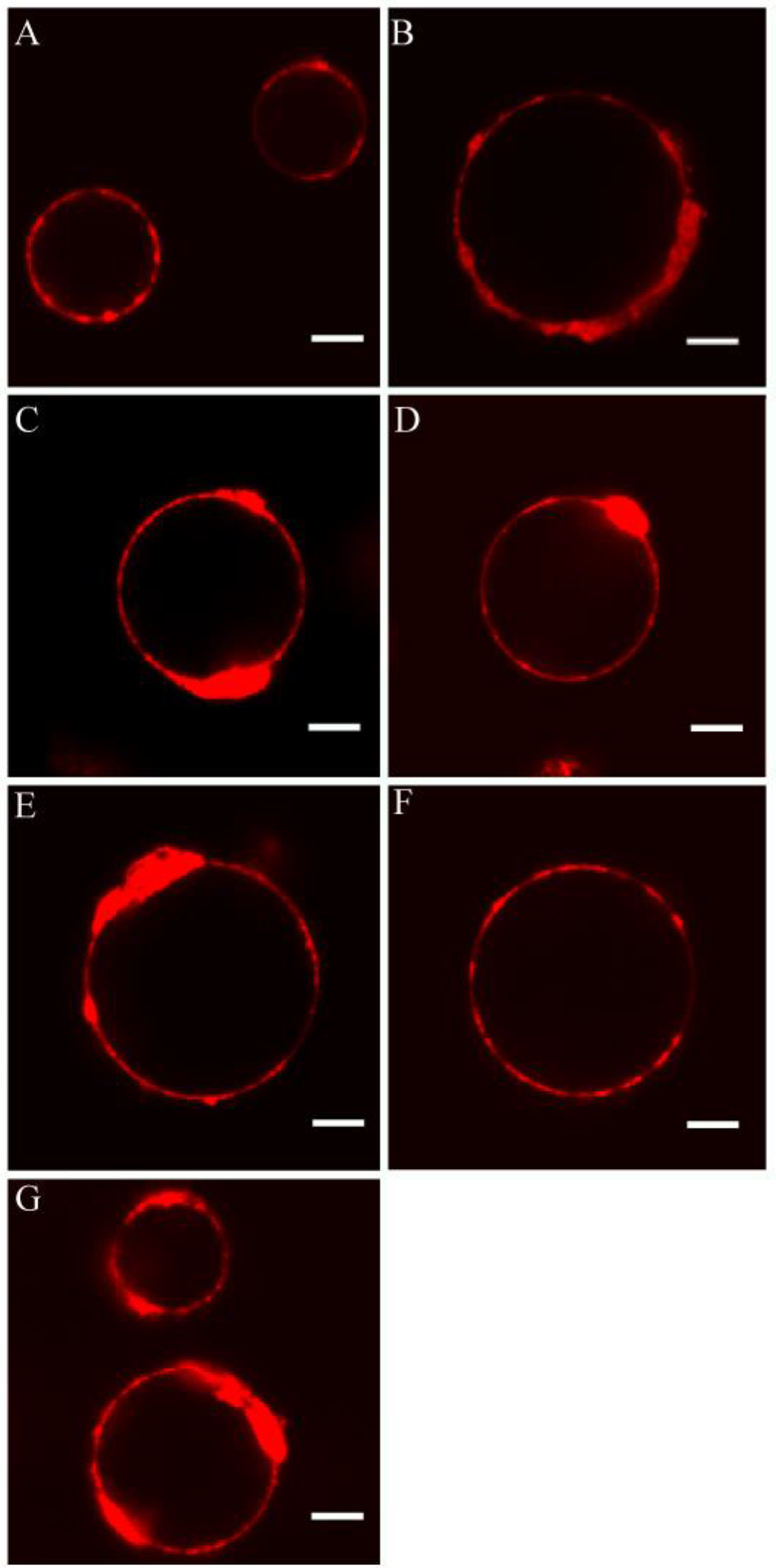
Chloroplast division mutants contain thylakoids that are much larger than those in wild-type plants. A-G) Representative cross-sections of osmotically swollen thylakoids from wild-type, *arc6, pdvl-1, pdvl-2, pdv2-1, pdv2-2*, and *arc3* plants, respectively, were taken using a laser scanning confocal microscope. Chlorophyll autofluorescence is used to track the presence of the thylakoid membrane. The scale bar corresponds to 10 μm.

The values in Table 4.1 report the average diameters of 1000 bleb samples. There was, however, a remarkable variation in the sizes of the blebs formed from these (and other) plants, and this is seen clearly in the diameter distribution histograms in Figure 5A. The possible origins of this variation are examined in the Discussion.

Together with the measurements of the electrical unit size above, the confocal microscopy images in Figures 3 and 4 show that *arc6* chloroplasts possess giant thylakoids. This suggests that thylakoid division is dependent on chloroplast division.

### Incomplete formation of contractile rings result in giant thylakoids

To test the dependence of thylakoid division on the constriction of the outer contractile ring, we examined two chloroplast division mutants, *pdv1* and *pdv2*, which exhibit 2-6 gigantic chloroplasts that possess a dumbbell-like structure. These proteins reside in the outer envelope membrane and act as a functional pair to interact with Arc 5 and form the outer contractile ring. The *pdv* mutant chloroplasts are still able to form the inner contractile ring, however without the outer ring the plastids never divide (Miyagishima et al., 2006). We tested two alleles of each *pdv* mutant, one that contained a mutation close to the N-terminus and another to the C-terminus which we labeled −1 and −2, respectively. On average, *pdv1-1, pdv1-2, pdv2-1*, and *pdv2-2* blebs (Figure 3C-F) were much larger than wild-type blebs. The blebs found in these mutants were slightly smaller than those found in the *arc6* mutant (~20 μm vs ~22 μm, Table 1). Representative z-projections show the *pdv* blebs in their entirety and provide further evidence that the blebs are much larger than those found in wild type (Figure 4C-F). The variations observed in bleb sizes for the wild type and *arc6* plants were evident in these mutants as well (Figure 5B).

**Table 1.** Bleb Dimensions. A table reporting the average bleb diameters and standard deviations for all genotypes. All values reported were statistically significant (P<0.05) from a student’s t-test other than those annotated with an (*). Samples annotated with (*) are not statistically significant from each other (P>0.05).

| Genotype | Diameter ($\mu\text{m}$) |
| --- | --- |
| wild type | $12.8 \pm 3.9$ |
| *arc6* | $21.7 \pm 11.2$ |
| *pdv1-1* * | $19.5 \pm 8.6$ |
| *pdv1-2* * | $20.1 \pm 8.8$ |
| *pdv2-1* * | $18.9 \pm 6.9$ |
| *pdv2-2* * | $19.8 \pm 7.8$ |
| *arc3* | $15.6 \pm 6.5$ |

*For all samples n= 1,000*

**Figure 4.**
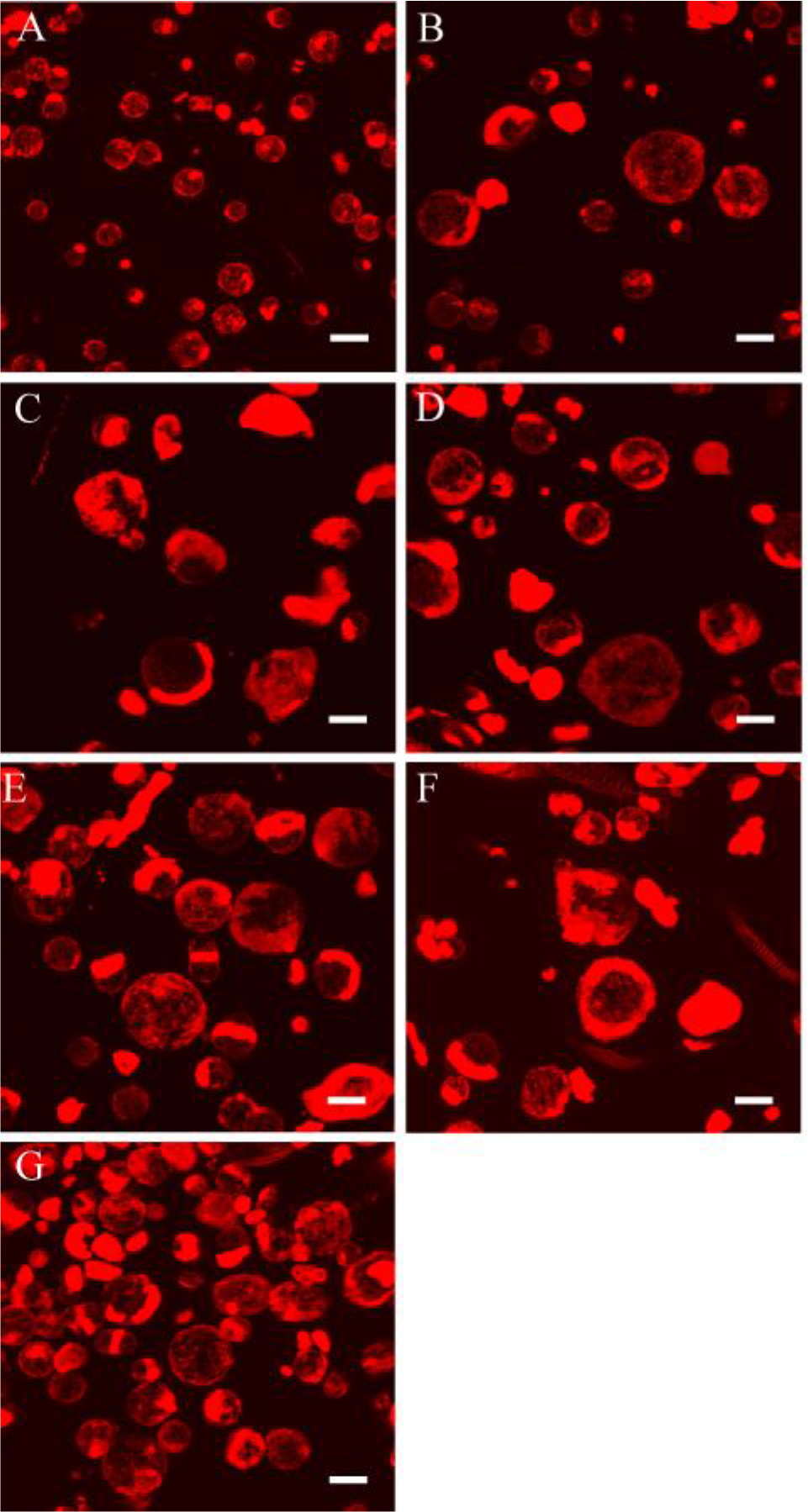
Z-projections of various thylakoid division mutants. Representative z-projections of osmotically swollen thylakoids from A) wild-type, B) *arc6*, C) *pdv1-1*, D) *pdv1-2*, E) *pdv2-1*, F) *pdv2-2*, and G) *arc3* plants were taken using a laser scanning confocal microscope. The scale bar corresponds to 20 μm.

To examine the behavior of thylakoids in a mutant that undergoes asymmetric plastid division we measured bleb sizes from an *arc3* mutant (Figure 3G) in which the inner contractile ring is misplaced (Pyke and Leech, 1994; Zhang et al., 2013). The *arc3* mutant plastids possess ~ 15 chloroplasts per cell (Burch-Smith et al., 2007). Here we expected the bleb sizes to average closer to those of the wild-type blebs, but with a larger variation, as an asymmetric division should result in one larger-than-wild-type and one smaller-than-wild-type chloroplast. As per these expectations, the *arc3* bleb sizes averaged 15.6 μm, somewhat larger than the wild-type blebs, and displayed a standard deviation of ± 6.5 μm (Table 1, Figure 5C). While we obtained a similar large standard deviation of bleb sizes from the giant chloroplast mutants, we note that blebs formed from the arc3 mutants are only 22% larger than those from wild type plants, but display a 67% increase in the diameter standard deviation.

**Figure 5.**
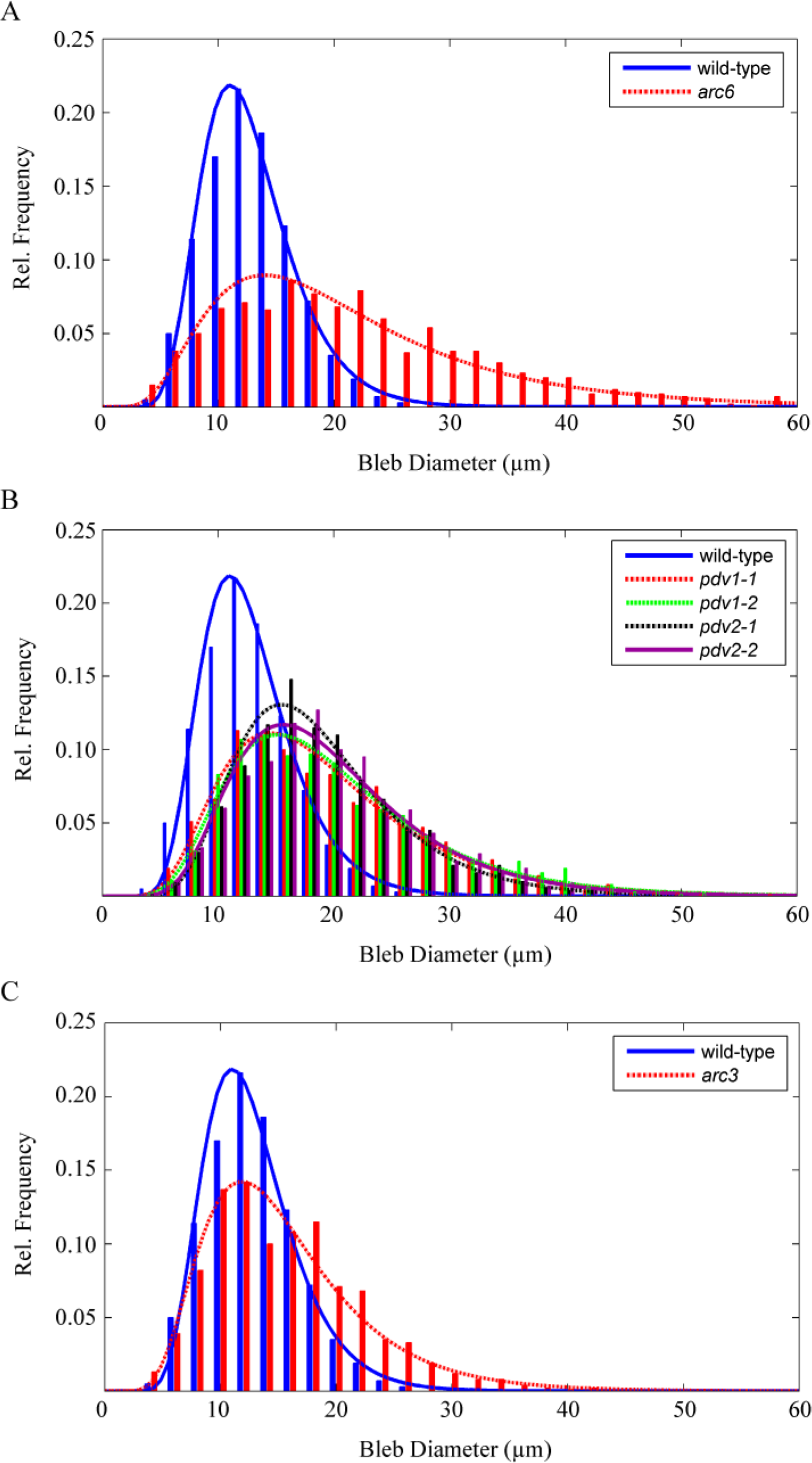
Thylakoid bleb population size distribution histogram. A) A histogram comparing the frequency of bleb diameter sizes between the wild type and A) *arc6*, B) all *pdv* mutants, and C) *arc3* plants (n=1,000 for all genotypes). The plots were fit with a log-normal distribution function.

These results in aggregate reveal that thylakoids do not possess an independent division machinery and cannot divide if the chloroplasts cannot divide.

## Discussion

### Thylakoid division is dependent on the chloroplast division machinery

Thylakoids represent a membrane system essential to shaping and maintaining the biosphere. Yet, the reproduction and partitioning of thylakoids, which must occur multiple times in green plants during each cell cycle, has so far escaped much scrutiny. In this work we have examined the question of whether thylakoids possess their own division machinery that operates independently of the chloroplast reproduction events.

In wild type plants it has been shown that each chloroplast possess a single thylakoid (Heslopharrison, 1963; Paolillo and Falk, 1966; Nierzwicki-Bauer et al., 1983; Schoenknecht et al., 1990; Shimoni et al., 2005; Mustardy et al., 2008; Austin and Staehelin, 2011; Daum and Kuhlbrandt, 2011). We reasoned that there are two possibilities for the structure of thylakoids found in plastid-division mutants. If the thylakoids are capable of dividing independently of chloroplast division, then the thylakoids in the plastid division mutants should be the same size as in wild-type chloroplasts. If, on the other hand, thylakoid division is dependent upon chloroplast division, then we would expect larger-than-wild-type thylakoids. We used two completely independent and unrelated methods to show that larger thylakoids are found in mutants that make giant chloroplasts. Changes in the conductivity of the thylakoid membrane in response to gramicidin addition can be used as a measure of relative thylakoid size (Schoenknecht et al., 1990). At any given ionophore concentration, it is expected that a larger vesicle would incorporate more gramicidin pores per vesicle than would a smaller one. Experimentally, this would be manifested as an increased sensitivity of the decay of the membrane potential to gramicidin. Using the amplitude of the carotenoid electrochromic signal at the end of a 9 ms illumination pulse as an indicator of the membrane conductivity, we found
that the *arc6* thylakoids were indeed considerably more sensitive to gramicidin addition in the manner suggested. From this we conclude that thylakoid vesicles in the *arc6* mutant are larger than those in the wild type. This would imply that the thylakoids cannot divide independently of plastid division.

It should also be noted that the ratio of gramicidin/chlorophyll required to accelerate the rate of electric field dissipation is higher in our studies than what was previously reported by (Schoenknecht et al., 1990). This discrepancy can be accounted for as a result of differences in thylakoid preparations. The thylakoids utilized in our studies were prepared by lysing intact chloroplasts within the same chamber that was carried out for taking the ΔA_520_ nm measurements. As a result, the thylakoid samples contained additional chloroplast envelope membranes which absorbed gramicidin and increased the working gramicidin/chlorophyll ratio up to 50 fold (Nishio and Whitmarsh, 1991). Thus the higher gramicidin concentrations used in this study are to be expected. Additionally, differences in gramicidin’s dimerization constants in thylakoids from different plant species can also contribute to this discrepancy (Schoenknecht et al., 1992). The difference in membrane permeability between the *arc6* and wild type thylakoids exhibited in Figure 2A-B may result from *arc6* thylakoids possessing naturally more ionically conductive membranes. It is also likely that the chloroplast solution used for the ECS measurements contained intact and broken chloroplasts which may have resulted in a slightly accelerated ΔA_520_ nm decay due to the damage that the thylakoids may have sustained during the event that caused the lysis of the intact chloroplast. However, the sensitivity of the thylakoids towards increasing concentrations of gramicidin would not be expected to be due to the presence of lysed chloroplasts. Accordingly, our observation that the *arc6* thylakoids are more sensitive towards increasing amounts of gramicidin indicates that the *arc6* thylakoids are larger than their wild type counter parts.

The sizes of osmotically lysed thylakoids from the different chloroplast division mutants provided an independent measurement of thylakoid structure. It is apparent that the thylakoids formed from *arc6* chloroplasts were considerably larger than wild-type blebs. The average diameter of the wild-type bleb was 12.8 μm, whereas that of *arc6* blebs was 21.7 μm (Table 1). Our similar findings with the *pdv1* and *pdv2* mutants suggests that thylakoid division is closely tied to the chloroplast division process (Table 1). The *arc6* chloroplasts are unable to form contractile rings, whereas the *pdv1* and *pdv2* mutants are capable of forming an inner contractile ring but only a partial outer ring. Our results suggest that separation of thylakoid membranes depends on the constriction forces applied by the chloroplast division machinery. The sizes of the *arc3* blebs show that thylakoid division is closely tied with the final stages of plastid division since the aberrant placement of the plastid ring leads to a rather large spread in thylakoid bleb sizes. Thus, our results from the bleb studies agree with the previously published electron microscopy images showing that thylakoid division occurs at the late stages of plastid division (Whatley, 1980; Robertson et al., 1996).

It is noteworthy that we observe a distribution of bleb sizes in our experiments (Figure 5), even in the wild type plants, the reasons for which may be manifold. First, the plastid division mutants we examined do not display giant chloroplasts in every cell type. Measurements of chloroplasts size found in the *arc6* guard cells made by (Pyke et al., 1994) and (Robertson et al., 1995) show that the *arc6* chloroplasts in the guard cells are smaller than the ones observed in mesophyll cells. Similarly, guard cells in *pdv* mutants also have smaller plastids compared to mesophyll cells (our observation). All cell types would have been in our preparations. Upon reflection one sees that a small number of cells with a normal allotment of chloroplasts mixed in with a preponderance of cells containing one and or a few giant chloroplasts would lead to a skewed bleb size distribution, as seen in our experiments. Second, it is clear from the patchiness of the observed chlorophyll fluorescence that thylakoids do not unstack and unravel completely during bleb formation, and incompletely expanded thylakoids would form smaller blebs. This incomplete unfolding was not a consequence of short incubations in water, as neither did the observed patchiness decrease (not shown), nor did the bleb diameters increase with longer incubation times (Figure S1). Third, while we made every effort to be gentle with the samples, we cannot be sure that we did not cause some breakage of the blebs during handling. We would expect osmotically swollen vesicles to be more fragile than normal thylakoids, and any such breakage would necessarily result in the formation of smaller blebs. Fourth, budding as an alternative plastid division mechanism has been observed in the tomato suffulta mutant (Chen et al., 2009), and *Bryophyllum pinatum* (Kulandaivelu and Gnanam, 1985). In a study that targeted GFP to the stroma of *arc6* mutant chloroplasts, small vesicular bodies that contained GFP were suggested to have budded off of the giant chloroplasts (Forth and Pyke, 2006). Such buds would be expected to contain smaller thylakoid vesicles. Finally, there is inherent variability in our samples. Wild type Arabidopsis cells do not always contain 100 chloroplasts, *arc6* mutant cells do not always contain one chloroplast, and *pdv1* mutant cells do not always contain three chloroplasts. Instead, these are averages. Thus we would expect some variability in bleb sizes even if the other factors mentioned above were not in play. We hold that the fact that we see any giant blebs at all in giant chloroplast mutants provides strong evidence that thylakoid division and chloroplast replication are not independent and uncoupled events.

### Thylakoid division mechanism

We propose that the constriction of the envelope membrane by the chloroplast division machinery acts to partition portions of the thylakoid into the two different poles of the dividing organelle. To reach this stage wherein a small portion of the thylakoid vesicle is trapped in the central isthmus of the dividing chloroplast, proteins that are known to be involved in thylakoid formation and remodeling, such as Vipp1, Thf1, FtsZ and FZL (Kroll et al., 2001; Wang et al., 2004; Gao et al., 2006; El-Kafafi et al., 2008; Lo and Theg, 2012), might be brought to bear on the thylakoid structure. These proteins have been proposed to govern the fission and fusion events accompanying stacking and unstacking of the thylakoid membrane in response to different light conditions (Chuartzman et al., 2008), and are thus candidates for those that might be involved in the final separation of the thylakoid vesicle into the two daughter plastids. Mesophyll cells in which the thylakoid protein FtsZ1, FtsZ2-2 and FZL has been knocked out possess fewer and larger chloroplasts compared to wild type cells (Gao et al., 2006; El-Kafafi et al., 2008; Karamoko et al., 2011), which suggests the involvement of thylakoid remodeling proteins during thylakoid division and also highlights the interdependence of both thylakoid and chloroplast division machineries for the complete fission of the daughter organelle. Our findings do not negate the possibility that other specific thylakoid division proteins are activated during a late stage of chloroplast division, but those proteins have yet to be identified. It appears more likely that the forces generated by the chloroplast envelope contractile rings are those that are responsible for dividing the thylakoid membrane as well. It will surely be interesting to elucidate through future studies the mechanism through which these forces generated on one membrane system are transduced to the other.

## Materials and Methods

### Plant material, protoplast and chloroplast isolation

The Col-0 ecotype of *Arabidopsis thaliana* was used as the wild type. The chloroplast division mutants, *arc6* (SAIL_693_G04), *pdv1-1, pdv1-2, pdv2-1*(SALK_059656), and *pdv2-2* (SAIL_875E10) were kindly provided by Dr. KW Osteryoung. Landsberg Erecta *arc3* (CS264) seeds were obtained from ABRC. All plants were genotyped and only homozygous lines were used (see supplemental materials for further detail). All plants were grown for 5 weeks on Murashige and Skoog (Phytotechnologies, Santa Cruz, CA) agar before harvesting the tissue for chloroplasts (Theg and Tom, 2011); growth chamber conditions were 20°C with 16 hrs light cycle of 100 μmol photons/m^2^/sec at 60% humidity. Plant tissue was harvested following the procedure described previously (Fitzpatrick and Keegstra, 2001) with some minor modifications. Briefly, plants were minced in a petri dish and washed with digestion buffer (400 mM sorbitol, 0.5mM CaCl_2_, 20 mM Mes-KOH, pH 5.2). The minced plant tissue was then incubated with 0.05 g/mL cellulase ‘onozuka’ R-10 and 0.01 g/mL macerozyme R-10 (Yakult Pharmaceutical Ind, Tokyo, Japan) for 3 hrs at room temperature with gentle rocking. Digested samples were passed through 4 layers of cheese cloth, and then centrifuged at 370 x *g* for 5 minutes at 4°C. The pellet was gently resuspended in resuspension buffer (400 mM sorbitol, 0.5mM CaCl_2_, 20 mM Mes-KOH, pH 6.0). Samples were centrifuged again at 660 x *g* for 5 minutes at 4°C, and if blebs were to be formed, the pellet was resuspended in a minimal volume of resuspension buffer and kept on ice in the dark until further use.

For wild-type chloroplast isolation, the protoplast pellet from the wild-type plants was resuspended in breakage buffer (330 mM Sorbitol, 5 mM EDTA, 5 mM EGTA, 10 mM NaHCO_3_, 0.1% BSA, 20 mM Tris-KOH, pH 8.4) and passed through 20 μm and 10 μm mesh through a syringe 4 times. For *arc6* chloroplast isolation, the protoplast pellet from the *arc6* tissues was resuspended in breakage buffer and incubated for 5 minutes on ice in the dark. Both samples were then passed through a Percoll gradient containing half volume of Percoll and half volume of grinding buffer (330 mM Sorbitol, 1 mM MgCl_2_, 1 mM MnCl_2_, 2 mM EDTA, 0.1% BSA, 50 mM Hepes-KOH, pH 7.3) at 8035 x *g* at 4°C. Intact chloroplasts were removed from the bottom band, and washed with a storage buffer (330 mM Sorbitol, 50 mM Hepes-KOH, pH 8). Chloroplasts were centrifuged at 1475 x *g* at 4°C for 5 min. This step was repeated again to remove the remaining Percoll. After chlorophyll determination (Arnon, 1949), isolated chloroplasts were stored in the storage buffer on ice in the dark until used.

### Electrochromic shift measurements

All electrochromic shift measurements contained 0.02 μg/μL chlorophyll concentration, 1 mM methyl viologen, and various amounts of gramicidin D (Sigma-Aldrich, St. Louis, Missouri); measurements were performed in 1.2 mL of storage buffer. Gramicidin stock solutions were prepared in ethanol; the total percentage of ethanol did not exceed 0.03% for all titrations. The measurements typically began with the addition of chloroplasts into a 2.5 ml polystyrene cuvette (Fisher Scientific, Houston, Texas) containing a master mix of methyl viologen, storage buffer and gramicidin. Samples were mixed with a stir bar for 2 minutes at room temperature, and then transferred into a 1.5 mL Polystyrene cuvette (Fisher Scientific, Houston, Texas) with a 10 mm path length. The absorbance readings were performed using a JTS-10 LED pump-probe spectrometer (Bio-Logic SAS, Claix, France). All 520 nm absorption measurements consisted of a 1 sec dark baseline followed by a 9 ms actinic pulse; the relaxation kinetics were followed out to 4 sec.

### Thylakoid bleb formation

Blebs were formed by diluting isolated protoplasts containing 2μg of chlorophyll into 2 mL of doubly distilled water stored at 4° C, and incubated on ice in the dark on ice for 1 hr before images were taken. Equal volumes of sample were loaded into a homemade perfusion chamber consisting of 1.8 mm x 100 mm x 1.1 mm piece of polycarbonate and a 50 x 22 mm coverslip (Fisher Scientific Houston, TX) which had been coated with poly-L-lysine (Sigma-Aldrich St. Louis, MO). Samples were centrifuged for 15 minutes at 60 x *g* using a GS-6KR swinging bucket centrifuge (Beckman Coulter Inc., Brea, CA) at 10°C and imaged.

### Microscopy

Phase contrast microscopy was performed with a Zeiss Standard 25 ICS microscope. Images at 160x magnification were taken by placing a camera, Canon powershot A620 (Canon, Melville, New York), into the ocular eyepiece. Fluorescence microscopy was performed using a Zeiss LSM 710 confocal microscope (Zeiss, Oberkochen, Germany). Image compilation and analysis were performed using Fiji software. Bleb diameters were measured from populations of isolated blebs taken from protoplast isolation preparations. The reported bleb diameter size were determined by scanning through the z-stack and finding the maximal diameter of the bleb. All reported values for bleb diameters were determined as being the largest distance from one end of the membrane to the other throughout the entire z-axis per bleb.

## Supplemental Material

**Figure S1.**
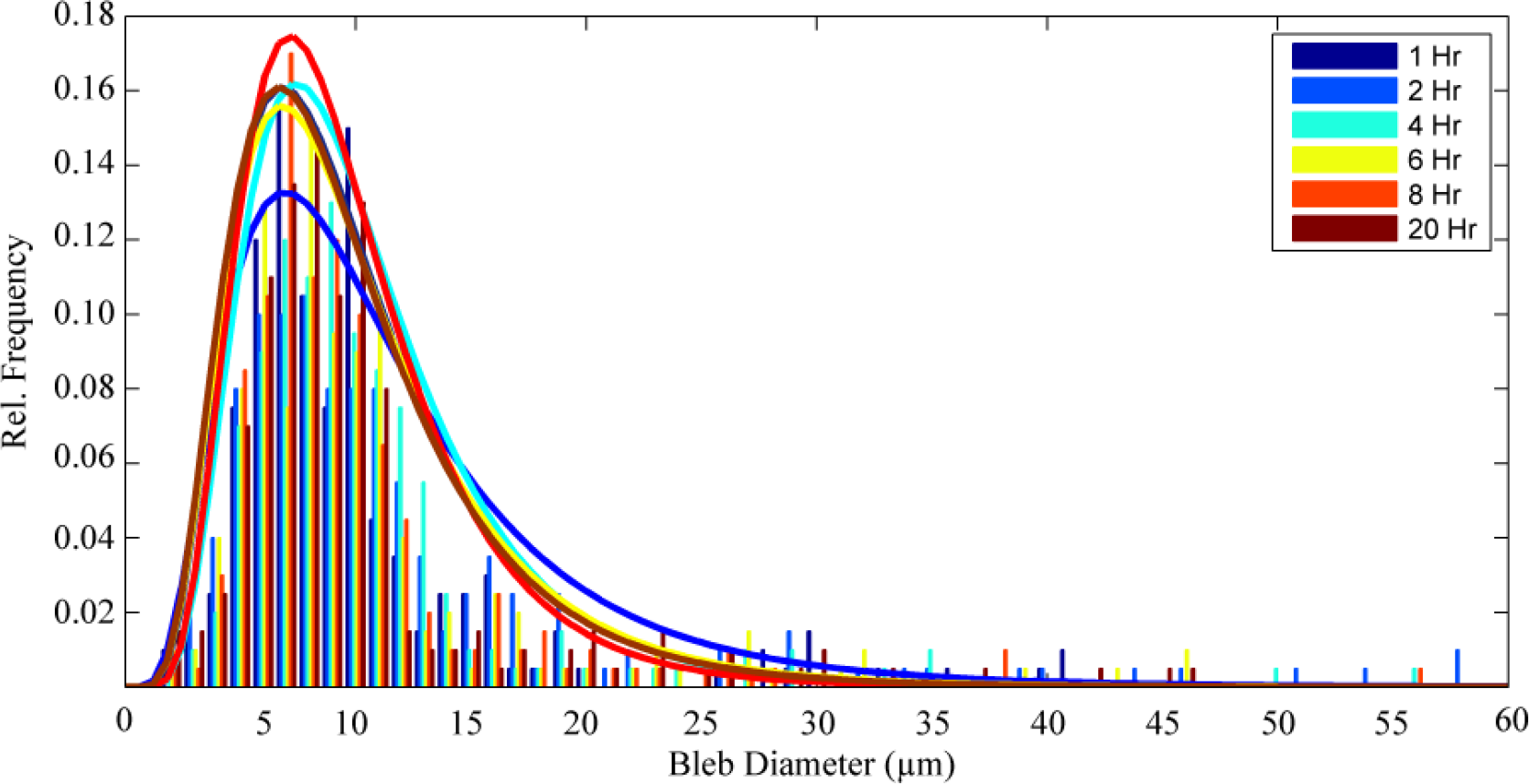
Thylakoid blebbing time course. A histogram reporting the frequency of bleb diameter sizes from a mixture of *arc6* and wild type samples over the course of 20 hours. (n=200 for all time points) The plots were fit with a log-normal distribution function.

**Table S1.**
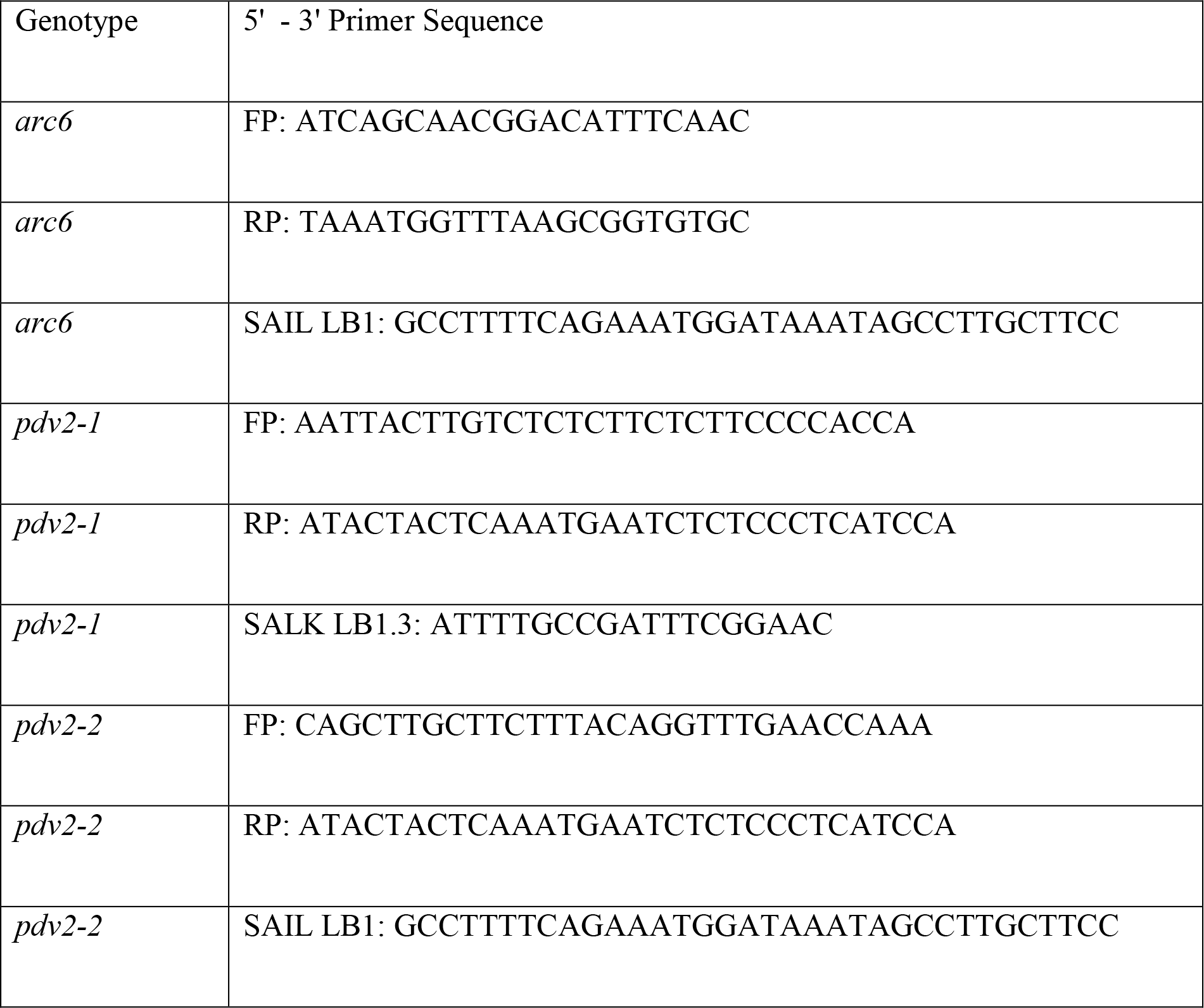
Table displaying the primers utilized to genotype plants from the T-DNA insertion lines, *arc6, and pdv2 −1, pdv2-2*. Plants from the T-DNA insertion lines, *arc6, and pdv2 −1, pdv2-2*, were genotyped using the following primers:

**Table S2.**
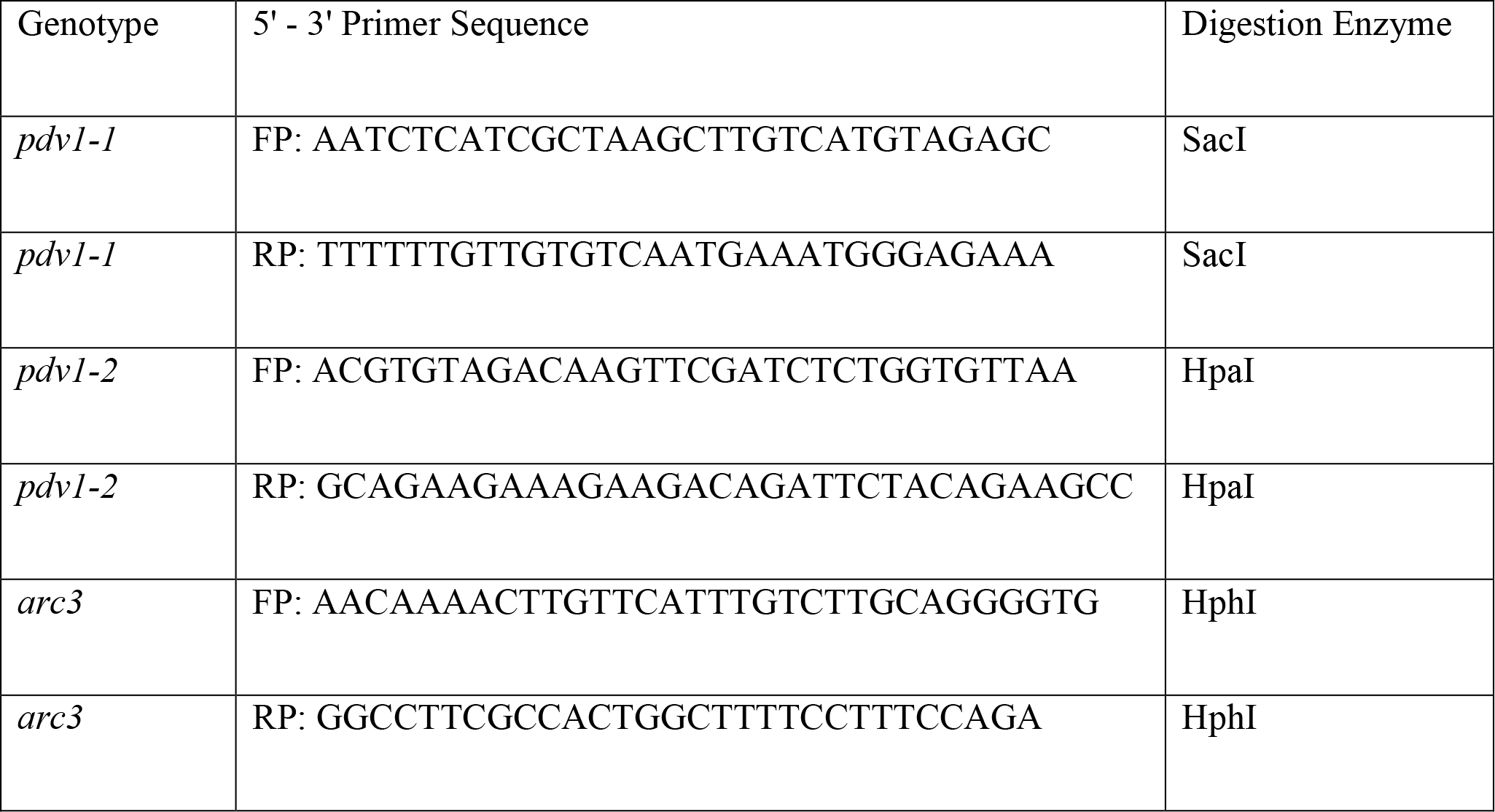
Table displaying the primers utilized to genotype *Pdv1-1, pdv1-I*, and *arc3* plants by detecting single nucleotide polymorphisms.

## Acknowledgements

We would like to thank Dr. Katherine W. Osteryoung for providing us with *arc6, pdv1-1, pdv1-2, pdv2-1*, and *pdv2-2 Arabidopsis* seeds, Drs. Bo Liu and Yuh-Ru Julie Lee for providing access to their phase contrast microscope and for sharing their expertise in confocal microscopy. We would also like to thank Drs. Lan-Xin Shi and Li Liu for their expertise in plant tissue culture. This work was funded by the Division of Chemical Sciences, Geosciences, and Biosciences, Office of Basic Energy Sciences of the US Department of Energy through Grant DE-FG02- 03ER15405.

## References

Arnon DI (1949) Copper Enzymes in Isolated Chloroplasts. Polyphenoloxidase in Beta Vulgaris. Plant Physiol 24: 1–15

Austin JR, 2nd, Staehelin LA (2011) Three-dimensional architecture of grana and stroma thylakoids of higher plants as determined by electron tomography. Plant Physiol 155: 1601–1611

Bailleul B, Cardol P, Breyton C, Finazzi G (2011) Electrochromism: a useful probe to study algal photosynthesis (vol 106, pg 179, 2010). Photosynthesis Research 110: 151–152

Boardman NK, Thorne SW, Anderson JM (1966) Fluorescence Properties of Particles Obtained by Digitonin Fragmentation of Spinach Chloroplasts. Proceedings of the National Academy of Sciences of the United States of America 56: 586–&

Boekema EJ, van Breemen JFL, van Roon H, Dekker JP (2000) Arrangement of photosystem II supercomplexes in crystalline macrodomains within the thylakoid membrane of green plant chloroplasts. Journal of Molecular Biology 301: 1123–1133

Burch-Smith TM, Schiff M, Caplan JL, Tsao J, Czymmek K, Dinesh-Kumar SP (2007) A novel role for the TIR domain in association with pathogen-derived elicitors. Plos Biology 5: 501–514

Chen YL, Asano T, Fujiwara MT, Yoshida S, Machida Y, Yoshioka Y (2009) Plant Cells Without Detectable Plastids are Generated in the crumpled leaf Mutant of Arabidopsis thaliana. Plant and Cell Physiology 50: 956–969

Chuartzman SG, Nevo R, Shimoni E, Charuvi D, Kiss V, Ohad I, Brumfeld V, Reich Z (2008) Thylakoid membrane remodeling during state transitions in Arabidopsis. Plant Cell 20: 1029–1039

Daum B, Kuhlbrandt W (2011) Electron tomography of plant thylakoid membranes. Journal of Experimental Botany 62: 2393–2402

Daum B, Nicastro D, Il JA, McIntosh JR, Kuhlbrandt W (2010) Arrangement of Photosystem II and ATP Synthase in Chloroplast Membranes of Spinach and Pea. Plant Cell 22: 1299–1312

El-Kafafi ES, Karamoko M, Pignot-Paintrand I, Grunwald D, Mandaron P, Lerbs-Mache S, Falconet D (2008) Developmentally regulated association of plastid division protein FtsZ1 with thylakoid membranes in Arabidopsis thaliana. Biochemical Journal 409: 87–94

Fitzpatrick LM, Keegstra K (2001) A method for isolating a high yield of Arabidopsis chloroplasts capable of efficient import of precursor proteins. Plant J 27: 59–65

Forth D, Pyke KA (2006) The suffulta mutation in tomato reveals a novel method of plastid replication during fruit ripening. J Exp Bot 57: 1971–1979

Gao H, Sage TL, Osteryoung KW (2006) FZL, an FZO-like protein in plants, is a determinant of thylakoid and chloroplast morphology. Proc Natl Acad Sci U S A 103: 6759–6764

Glynn JM, Froehlich JE, Osteryoung KW (2008) Arabidopsis ARC6 coordinates the division machineries of the inner and outer chloroplast membranes through interaction with PDV2 in the intermembrane space. Plant Cell 20: 2460–2470

Heslopharrison J (1963) Structure and morphogenesis of lamellar systems in grana-containing chloroplasts. Planta 60: 243–260

Heslopharrison J (1963) Structure and Morphogenesis of Lamellar Systems in Grana-Containing Chloroplasts .1. Membrane Structure and Lamellar Architecture. Planta 60: 243–260

Hinnah SC, Wagner R (1998) Thylakoid membranes contain a high-conductance channel. European Journal of Biochemistry 253: 606–613

Junge W, Witt HT (1968) On Ion Transport System of Photosynthesis - Investigations on a Molecular Level. Zeitschrift Fur Naturforschung Part B-Chemie Biochemie Biophysik Biologie Und Verwandten Gebiete B 23: 244–&

Karamoko M, El-Kafafi ES, Mandaron P, Lerbs-Mache S, Falconet D (2011) Multiple FtsZ2 isoforms involved in chloroplast division and biogenesis are developmentally associated with thylakoid membranes in Arabidopsis. Febs Letters 585: 1203–1208

Kroll D, Meierhoff K, Bechtold N, Kinoshita M, Westphal S, Vothknecht UC, Soll J, Westhoff P (2001) VIPP1, a nuclear gene of Arabidopsis thaliana essential for thylakoid membrane formation. Proc Natl Acad Sci U S A 98: 4238–4242

Kulandaivelu G, Gnanam A (1985) Scanning Electron-Microscopic Evidence for a Budding Mode of Chloroplast Multiplication in Higher-Plants. Physiologia Plantarum 63: 299–302

Leech RM, Thomson WW, Plattaloia KA (1981) Observations on the Mechanism of Chloroplast Division in Higher-Plants. New Phytologist 87: 1–&

Lo SM, Theg SM (2012) Role of Vesicle-Inducing Protein in Plastids 1 in cpTat transport at the thylakoid. Plant J

Maple-Grodem J, Raynaud C (2014) Plastid Division. In SM Theg, FA Wollman, eds, Plastid Biology. Springer, New York, pp 155–188

McAndrew RS, Froehlich JE, Vitha S, Stokes KD, Osteryoung KW (2001) Colocalization of plastid division proteins in the chloroplast stromal compartment establishes a new functional relationship between FtsZ1 and FtsZ2 in higher plants. Plant Physiology 127: 1656–1666

Mercer FV, Hodge, A.J., Hope, A.B., Mclean, J. D. (1954) The Structure and Swelling Properties of Nitella Chloroplasts. Australian Journal of Biological Sciences 8: 1–18

Miyagishima SY, Froehlich JE, Osteryoung KW (2006) PDV1 and PDV2 mediate recruitment of the dynamin-related protein ARC5 to the plastid division site. Plant Cell 18: 2517–2530

Mustardy L, Buttle K, Steinbach G, Garab G (2008) The Three-Dimensional Network of the Thylakoid Membranes in Plants: Quasihelical Model of the Granum-Stroma Assembly. Plant Cell 20: 2552–2557

Mustardy LA, Janossy AGS (1979) Evidence of helical thylakoid arrangement by scanning electron microscopy. Plant Sci. Lett. 16: 281–284

Nierzwicki-Bauer SA, Balkwill DL, Stevens SE, Jr. (1983) Three-dimensional ultrastructure of a unicellular cyanobacterium. J Cell Biol 97: 713–722

Nishio JN, Whitmarsh J (1991) Dissipation of the Proton Electrochemical Potential in Intact and Lysed Chloroplasts .1. The Electrical Potential. Plant Physiology 95: 522–528

Oross JW, Possingham JV (1989) Ultrastructural Features of the Constricted Region of Dividing Plastids. Protoplasma 150: 131–138

Osteryoung KW, Pyke KA (2014) Division and dynamic morphology of plastids. Annu Rev Plant Biol 65: 443–472

Paolillo D, Falk R (1966) The ultrastructure of grana in mesolhyll plastids of Zea mays. Am J Botany 53: 173–180

Pyke KA, Leech RM (1994) A Genetic-Analysis of Chloroplast Division and Expansion in Arabidopsis-Thaliana. Plant Physiology 104: 201–207

Pyke KA, Rutherford SM, Robertson EJ, Leech RM (1994) Arc6, a Fertile Arabidopsis Mutant with Only 2 Mesophyll Cell Chloroplasts. Plant Physiology 106: 1169–1177

Robertson EJ, Pyke KA, Leech RM (1995) Arc6, an Extreme Chloroplast Division Mutant of Arabidopsis Also Alters Proplastid Proliferation and Morphology in Shoot and Root Apices. Journal of Cell Science 108: 2937–2944

Robertson EJ, Rutherford SM, Leech RM (1996) Characterization of chloroplast division using the Arabidopsis mutant arc5. Plant Physiology 112: 149–159

Schoenknecht G, Althoff G, Junge W (1990) The electric unit size of thylakoid membranes. FEBS Lett. 277:65–68

Schoenknecht G, Althoff G, Junge W (1992) Dimerization constant and single-channel conductance of gramicidin in thylakoid membranes. Journal of Membrane Biology 126: 265–275

Shimoni E, Rav-Hon O, Ohad I, Brumfeld V, Reich Z (2005) Three-dimensional organization of higher-plant chloroplast thylakoid membranes revealed by electron tomography. Plant Cell 17: 2580–2586

Theg SM, Tom C (2011) Measurement of the DeltapH and electric field developed across Arabidopsis thylakoids in the light. Methods Mol Biol 775: 327–341

van Roon H, van Breemen JFL, de Weerd FL, Dekker JP, Boekema EJ (2000) Solubilization of green plant thylakoid membranes with n-dodecyl-alpha,D-maltoside. Implications for the structural organization of the Photosystem II, Photosystem I, ATP synthase and cytochrome b(6)f complexes. Photosynthesis Research 64: 155–166

Vitha S, Froehlich JE, Koksharova O, Pyke KA, van Erp H, Osteryoung KW (2003) ARC6 is a J-domain plastid division protein and an evolutionary descendant of the cyanobacterial cell division protein Ftn2. Plant Cell 15: 1918–1933

Wang Q, Sullivan RW, Kight A, Henry RL, Huang JR, Jones AM, Korth KL (2004) Deletion of the chloroplast-localized Thylakoid formation1 gene product in Arabidopsis leads to deficient thylakoid formation and variegated leaves. Plant Physiology 136: 3594–3604

Weier TE, Stocking CR, Bracker CE, Risley EB (1965) Structural Relationships of Internal Membrane Systems of in Situ and Isolated Chloroplasts of Hordeum Vulgare. American Journal of Botany 52: 339–&

Whatley JM (1980) Plastid Growth and Division in Phaseolus-Vulgaris. New Phytologist 86: 1–&

Witt HT (1979) Energy-Conversion in the Functional Membrane of Photosynthesis - Analysis by Light-Pulse and Electric Pulse Methods - Central Role of the Electric-Field. Biochimica Et Biophysica Acta 505: 355–427

Yoshida Y, Kuroiwa H, Misumi O, Nishida K, Yagisawa F, Fujiwara T, Nanamiya H, Kawamura F, Kuroiwa T (2006) Isolated chloroplast division machinery can actively constrict after stretching. Science 313: 1435–1438

Zhang M, Schmitz AJ, Kadirjan-Kalbach DK, Terbush AD, Osteryoung KW (2013) Chloroplast division protein ARC3 regulates chloroplast FtsZ-ring assembly and positioning in arabidopsis through interaction with FtsZ2. Plant Cell 25: 1787–1802

